# 3D-printing-assisted, microfabricated devices reveal hierarchical and temporal mechanosensing in high-density fibroblast culture

**DOI:** 10.1101/2025.09.18.677242

**Authors:** Ghiska Ramahdita, Xiangjun Peng, Mohammad Jafari, Charlene Pobee, Riya Bhakta, Zhuangyu Zhang, Missy Pear, Fei Wang, David Schuftan, Yuan Hong, Michael David, Elliot L. Elson, Chao Zhou, Guy M. Genin, Farid Alisafaei, Nathaniel Huebsch

## Abstract

Understanding how cells integrate mechanical forces across multiple directions, length scales, and timescales remains a fundamental challenge in mechanobiology. Deciphering how cells integrate this information is particularly important in the context of wound healing, where the timing and duration of the fibroblast-to-myofibroblast transition can determine healing outcomes. Here, we discovered that fibroblasts in engineered tissues respond to directional anisotropy in stress through a hierarchical temporal cascade, with individual cell elongation (24 hr) preceding collective alignment (40 hr), which then drives α-smooth muscle actin expression and myofibroblast transition (96h). To enable this discovery, we developed a modified hydrogel-assisted stereolithographic elastomer (HASTE) prototyping platform to incorporate a detergent that improves wettability of template agar hydrogels by poly(dimethylsiloxane) elastomer. HASTE allowed rapid prototyping of intricate 3D micropost arrays that provides isotropic (8-post) versus anisotropic (4-post) boundary conditions. Fibroblasts sensed and responded to stress directionality before bulk tissue reorganization occurs. Computational modeling predicted steady-state activation patterns based on initial stress anisotropy rather than magnitude, and our experiments reveal that reaching this state requires sequential mechanosensitive processes operating across distinct timescales. This temporal hierarchy persists even when extensive cell-cell contacts might be expected to mask matrix-mediated mechanical signals. Our findings demonstrate that fibroblast mechanosensing involves adaptive responses encoded through progressive cell and tissue reorganization. Results provide insight into how nanoscale mechanosensing scales up to direct tissue-level organization, with implications for understanding wound healing, understanding fibrosis, and engineering functional tissue replacements.

## 1. Introduction

A fundamental question in mechanobiology is not just what mechanical signals cells sense, but how they integrate these signals across time and length scales to make fate decisions. Living cells exist in dynamic environments where the spatiotemporal path to mechanical equilibrium may be as important as the final state itself. This spatiotemporal aspect of mechanosensing remains poorly understood, particularly in three-dimensional tissues where cells must coordinate their responses while simultaneously reshaping their mechanical environment.^1^

Fibroblast biology exemplifies this challenge. During wound healing and tissue repair, fibroblasts integrate mechanical cues from their environment when undergoing the transition to myofibroblasts, characterized by increased contractility and α-smooth muscle actin (α-SMA) expression.^2,3^ While this fibroblast-to-myofibroblast transition (FMT) aids wound closure,^4^ its dysregulation leads to fibrosis.^2,5–7^ A substantial body of literature has established that the physical properties of the ECM, especially matrix elasticity, and overall tissue tension are key mechanical signals driving FMT.^4,8–11^ However, these outcomes may also depend on the timing of the FMT, which is inducible over short experimental and wound-healing timescales and can persist chronically. In 2D culture, TGF-β1 or mechanical strain can induce proto-myofibroblast features within hours and detectable α-SMA transcription/protein incorporation and increased contractility within about 1-3 days, with maturation of stress fibers and force generation progressing over ∼3-7 days under continued stimulation.^8,12,13^ In wound repairmyofibroblasts typically emerge in granulation tissue over several days after injury and are prominent through the first 1-2 weeks, then regress if healing resolves; persistence beyond these windows is associated with fibrosis and scar formation.^3,4,14^ These timelines reflect the integrated effects of TGF-β signaling, mechanotransduction, and matrix context; sustained cues prolong the myofibroblast state, whereas their withdrawal leads to deactivation/apoptosis and loss of α-SMA stress fibers.^4,15–17^

However, these studies typically focus on scalar metrics of mechanical input, such as ECM stiffness or tension magnitude, without accounting for the directionality of stress. Even in their quiescent, unactivated state, fibroblasts are elongated cells.^18^ Recent studies have revealed that fibroblasts respond not only to the magnitude of mechanical stress but also to its directionality or anisotropy.^19^ The temporal sequences that lead to this mechano-responsiveness have not been identified, and these responses have not been studied in tissues with cell densities approaching the density of many natural tissues (e.g. >10^7^ cells/mL), where cell-cell contacts compete with cell-matrix interactions. In such high-density tissues, collective contractility, matrix compaction, and cell-cell interactions could override local mechanical cues, potentially blunting the effect of tensional anisotropy. Alternatively, it is possible that cells retain an intrinsic sensitivity to stress directionality, and that this sensitivity continues to regulate phenotypic transitions even in the setting of large-scale tissue reorganization.

To address the possibility that fibroblasts’ ability to sense tensional anisotropy depends on cell density, we developed a microfabricated platform capable of imposing either isotropic or anisotropic stress fields within tissues. Using a rapid prototyping approach based on stereolithographic printing and hydrogel-assisted molding,^20^ we created arrays of deformable microposts designed to impose either isotropic or anisotropic boundary conditions on fibroblast-laden collagen matrices. These devices allow for precise geometric control of the tissue stress field. By varying cell density and culture time, and integrating experimental observations with computational modeling, we examined how cells detect and respond to anisotropic mechanical environments under conditions that mimic real tissue architecture.

Our results reveal that tensional anisotropy governs a distinct sequence of events during fibroblast activation: early changes in cell elongation and alignment precede tissue-scale compaction and culminate in the upregulation of contractile proteins associated with myofibroblast transition. These effects are robust across a range of cell densities and persist even in the presence of high levels of cell-cell contact and matrix remodeling. Together, these findings identify stress anisotropy as a critical regulator of fibroblast fate in 3D tissues and offer a framework for understanding how geometry and mechanics together orchestrate fibroblast behavior during repair and disease.

## 2. Results

### Triton X-100 modification enables high-fidelity fabrication of complex 3D micropost arrays

The study of how mechanical boundary conditions influence cellular behavior requires precise control over tissue geometry and mechanical constraints. While microfabricated tissue gauges (“micro-TUGs”) have proven valuable for measuring cell-generated forces,^21^ their fabrication traditionally requires costly photolithography with limited design flexibility. To overcome this limitation, we adapted the recently developed HASTE (Hydrogel-Assisted Stereolithographic Elastomer) technique^20^ to create poly(dimethylsiloxane) (PDMS) microposts analogous to those first described by Legant, *et al.*^21^ (**Fig. 1a**).

**Figure 1.**
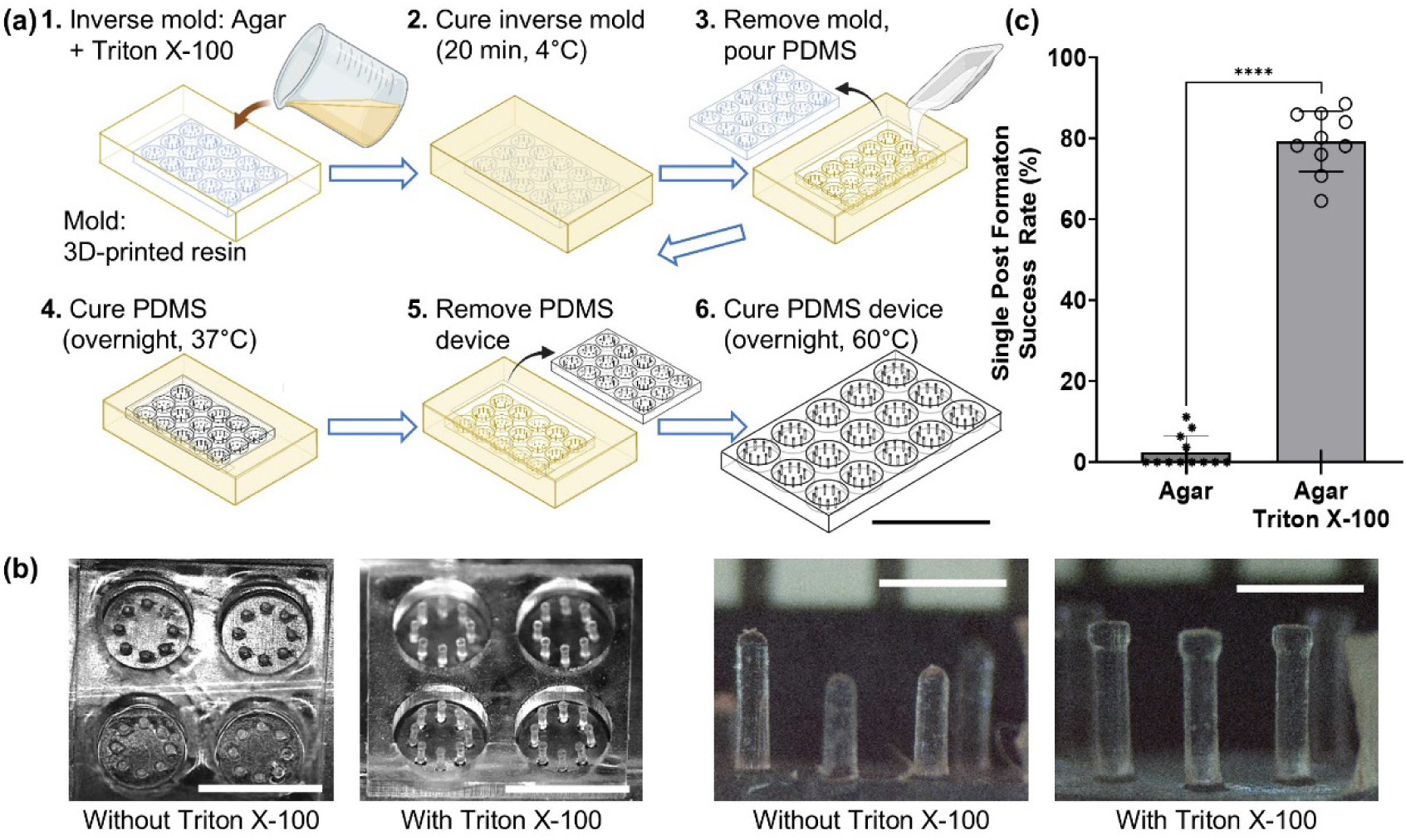
Surfactant-modified hydrogel molding enables reliable fabrication of high aspect-ratio microposts with overhanging caps. **(a)** Schematic of the HASTE (Hydrogel-Assisted Stereolithographic Elastomer) fabrication process. 3D-printed masters serve as positive molds to create agar hydrogel negative molds, which are then used to cast PDMS devices. The addition of 0.2% Triton X-100 to the agar (orange) reduces interfacial tension with hydrophobic PDMS, enabling complete infiltration into narrow mold features. **(b)** Representative brightfield images comparing micropost fabrication outcomes. Without Triton X-100, PDMS fails to completely fill the mold cavities, resulting in malformed or missing posts with incomplete caps. With 0.2% Triton X-100, uniform arrays of well-formed microposts with intact overhanging caps (300 μm diameter) are consistently achieved. Scale bars: left, 5 mm; right, 100 µm. **(c)** Quantification of fabrication success rate across multiple batches (*n* = 10 sets of post-based devices, with at least 96 individual posts per device batch, from four different batches of PDMS). Triton X-100 modification increased successful device yield (*p* < 0.0001).

HASTE uses 3D-printed masters and hydrogel intermediate molds, enabling rapid design iteration and cost-effective prototyping of complex geometries. However, fabricating high aspect-ratio microposts with overhanging caps - essential for preventing tissue detachment during contraction^21^ - presented a critical technical challenge. The hydrophobic PDMS prepolymer poorly infiltrated the narrow channels in hydrogel molds, resulting in incomplete features and frequent fabrication failures. We hypothesized that this occurred because of the interfacial energy mismatch between the hydrophilic agar and hydrophobic PDMS, and that adding a detergent to the water within the agar would lower this interfacial energy, improving overall wetting of the PDMS into the agar negative. Thus, we incorporated the mild detergent, Triton X-100 into the agar hydrogel at a concentration of 0.2%. This dramatically reduced interfacial tension between the hydrophobic PDMS and hydrogel during crosslinking, markedly enhancing the success rate of making devices: while standard agar molds yielded poorly formed or incomplete microposts, Triton X-100-modified molds consistently produced well-defined arrays with intact overhanging caps (**Fig. 1b,c**).

The flexibility of stereolithographic 3D printing, combined with our improved molding protocol, enabled facile control over both tissue geometry and microwell design. This capability allowed us to investigate how different mechanical boundary conditions, specifically isotropic (8-post) versus anisotropic (4-post) configurations, influence fibroblast behavior and fate decisions in 3D tissues.

### Engineered microtissues recapitulate fibroblast-mediated collagen compaction and remodeling

To investigate how mechanical boundary conditions influence fibroblast-to-myofibroblast transition, we developed a 3D culture system that captures the dynamic interplay between cells and their extracellular matrix during tissue remodeling. In wound healing, fibroblasts generate contractile forces that reorganize collagen, creating a mechanical feedback loop that drives phenotypic changes including cellular polarization and α-SMA upregulation.^22–24^ Our micropost platform was designed to impose controlled mechanical constraints while allowing cells to freely remodel their local environment.

When NIH/3T3 fibroblasts were embedded in type I collagen gels (1.5 mg/mL) within our microwells, they immediately began contracting and reorganizing the matrix through actomyosin-generated forces (**Fig. 2a**). This cell-mediated compaction progressively increased matrix density and collagen fiber alignment, particularly in regions of high cell density where enhanced cell-cell mechanical communication amplified collective contractility.^25,26^ Importantly, the compliant microposts (elastic modulus ∼130 kPa) deformed inward as tissues contracted, providing both mechanical resistance and a visual readout of tissue-generated forces.

**Figure 2.**
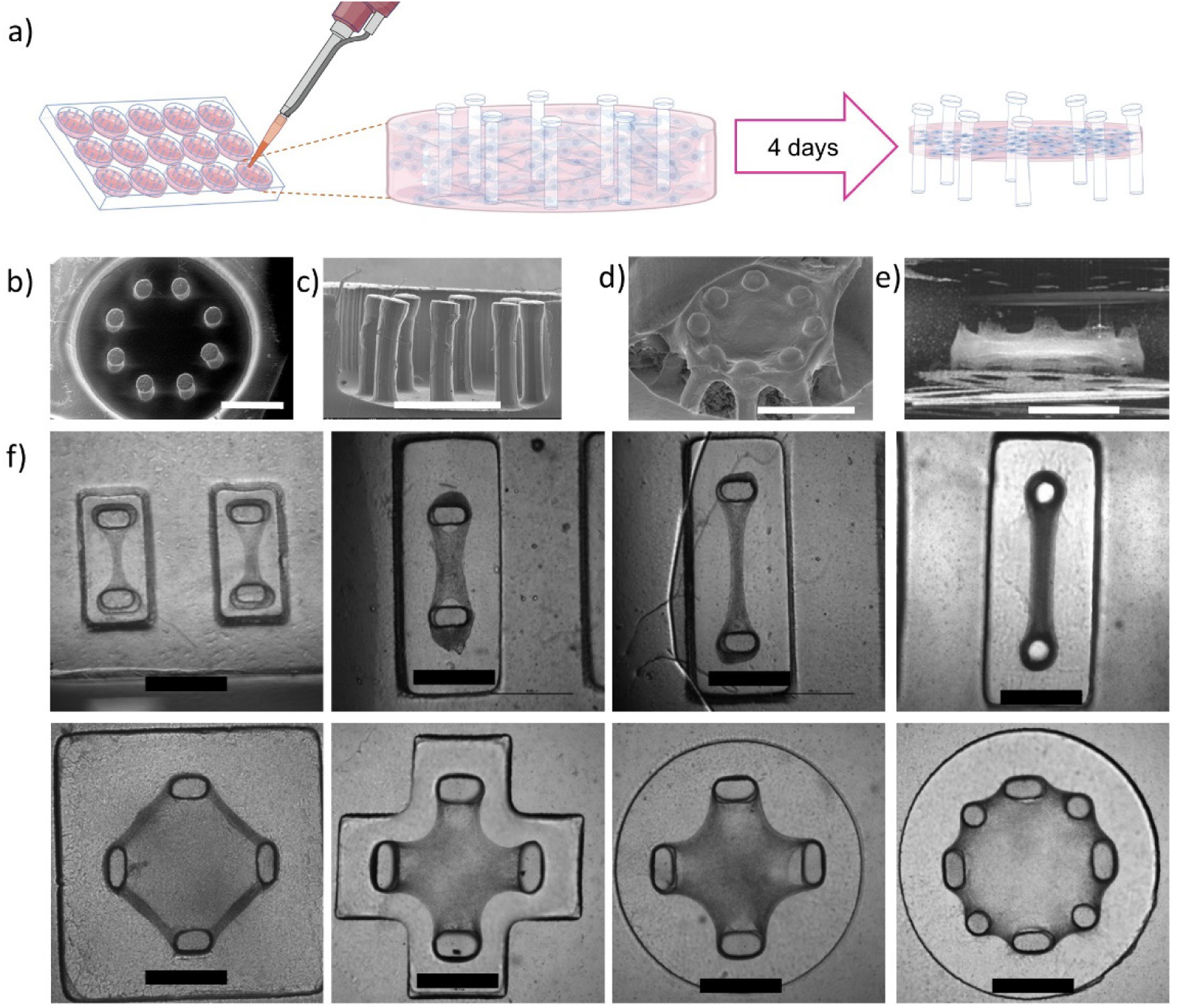
Fibroblast-laden collagen microtissues undergo cell-mediated compaction while suspended between compliant microposts. **(a)** Schematic illustrating progressive tissue remodeling over 4 days of culture. Fibroblasts embedded in collagen (1.5 mg/mL) generate contractile forces that compact the matrix, causing inward deflection of compliant PDMS microposts (130 kPa elastic modulus). Tissues detach from well surfaces within 24 hours, remaining anchored only to overhanging post caps (created with BioRender). **(b-d)** Environmental scanning electron microscopy (E-SEM) validation of tissue suspension and post-engagement after 4 days of culture. **(b)** Top view showing compacted tissue spanning between 8 posts with characteristic morphology. **(c)** Side view. **(d)** 45° oblique view revealing tissue thickness and three-dimensional organization around post caps. **(e)** Side-view of Optical coherence tomography (OCT) three-dimensional reconstruction confirming tissue suspension and quantifying tissue thickness (100-200 μm) and vertical position within the well. Microposts appear as vertical columns with tissue bridges visible between posts. **(f)** Design versatility enabled by HASTE fabrication platform. Representative examples of different micropost configurations including 4-post (anisotropic stress field), 8-post (isotropic center/anisotropic periphery), and custom geometries for specific mechanical boundary conditions. Each design maintains identical well diameter (3.26 mm) and post dimensions (200 μm diameter, 1.125 mm height) while varying spatial arrangement. Scale bars: 1 mm.

Environmental scanning electron microscopy (E-SEM) revealed that tissues completely detached from the bottom and sides of the PDMS wells within 24 hours, remaining anchored only to the overhanging caps of the microposts (**Fig. 2b-d**). This suspended configuration, confirmed by optical coherence tomography (OCT) imaging (**Fig. 2e**), ensures that mechanical signals are transmitted primarily through the tissue itself rather than through adhesion to rigid substrates. The tissues formed continuous structures spanning between posts, with the degree of compaction and post deflection increasing over the 4-day culture period.

The versatility of our fabrication approach enabled systematic exploration of how different geometric constraints influence tissue mechanics. By varying the number and arrangement of posts, we could impose either isotropic (8-post) or anisotropic (4-post) boundary conditions while maintaining identical culture conditions and initial tissue volumes (**Fig. 2f**). This capability to rapidly prototype different geometries proved essential for dissecting how stress field directionality influences fibroblast behavior, as explored in subsequent experiments.

### Computational modeling predicts that stress anisotropy, not stress magnitude, governs fibroblast activation

To understand how mechanical cues regulate fibroblast morphology and activation within engineered tissues, we used a multiscale continuum model to simulate the stress fields and predicted cell activation states in two geometrically distinct microtissue configurations: one defined by four boundary posts (4-post) and the other by eight (8-post). In this model, tissues are treated as a continuum of representative volume elements (RVEs), each of which is composed of an active contractile cell and collagen matrices (Materials and Methods, Supplementary Information).^19,27,28^ Starting with the same initially round shape in our circular micro-wells, our simulations first show the evolution of tissue shape in the 4-post and 8-post tissue designs (**Fig. 3a**). We confirmed experimentally that our micro-tissues compact in the manner predicted by the model (**Fig. 4**).

**Figure 3.**
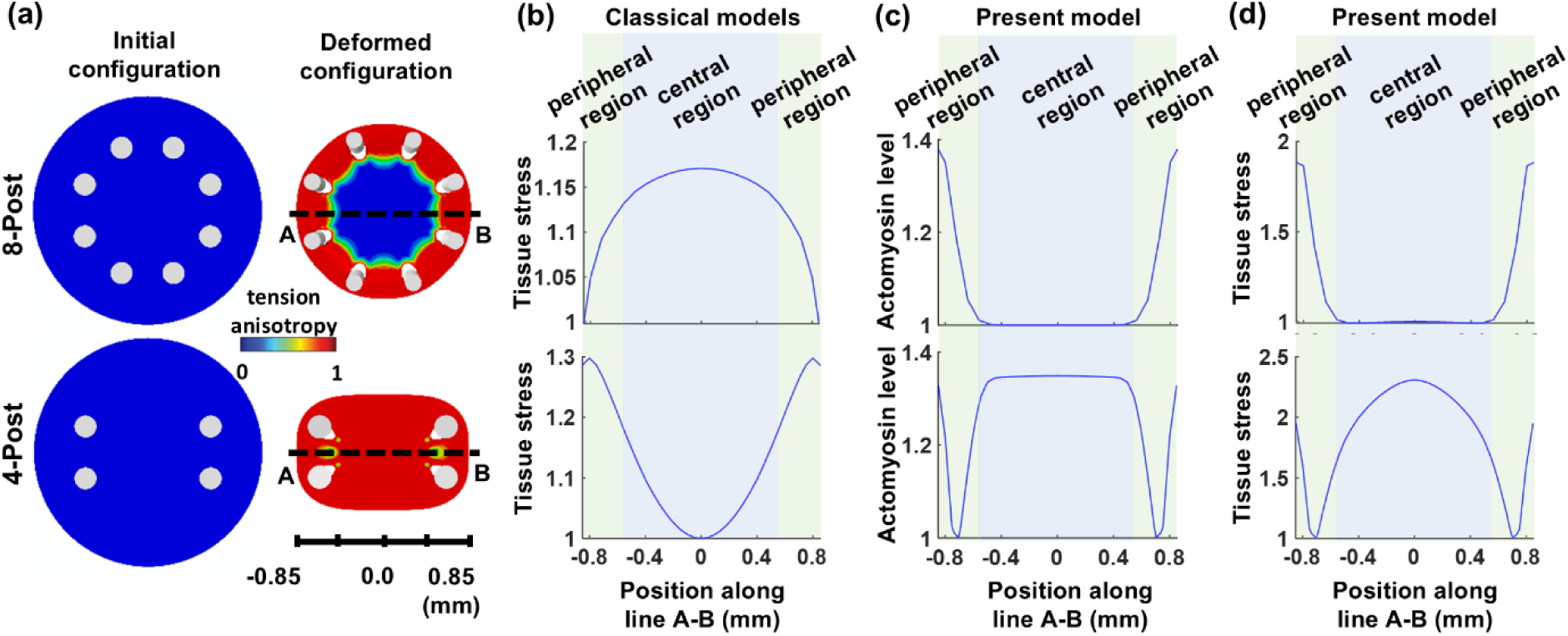
Computational modeling reveals opposing predictions between classical and anisotropy-driven activation models. **(a)** Simulated tissue compaction and stress field evolution in 8-post (top) and 4-post (bottom) configurations. Color maps indicate stress anisotropy (tanh[(𝜎₁/𝜎₂ − 1)/2]). The 8-post geometry generates an isotropic stress field at the tissue center (blue) with circumferential anisotropy at the periphery (red); the 4-post geometry produces predominantly anisotropic stress throughout, with maximum anisotropy along the horizontal axis between opposing posts. **(b)** Classical models predict the highest tension magnitude at the center of 8-post tissues and the minimum tension at the center of 4-post tissues. **(c)** The anisotropy-sensing model predicts the opposite activation pattern for actomyosin level, and **(d)** tissue stress, demonstrating how cellular mechanosensing of stress directionality can override effects of stress magnitude. Plots in **(b-d)** are normalized.

**Figure 4.**
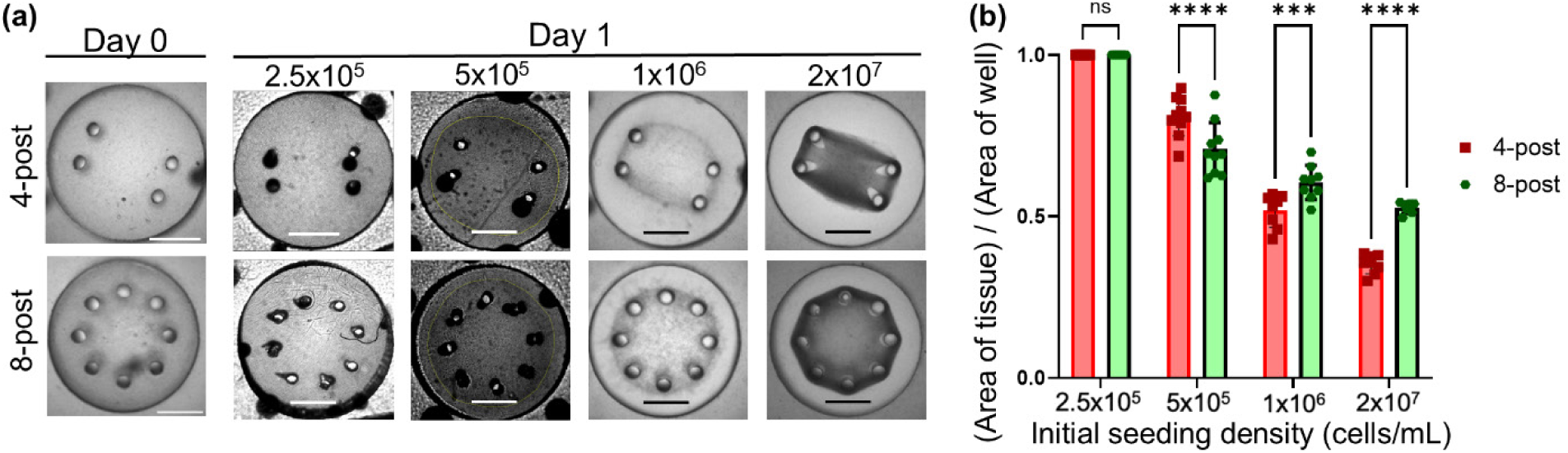
Cell density-dependent tissue compaction validates computational predictions of geometry-controlled mechanics. **(a)** Representative brightfield images demonstrating differential tissue compaction between 4-post and 8-post boundary conditions after 24 hours of culture. Tissues were seeded at four initial cell densities (0.25, 0.5, 1.0, and 20 × 10^6^ cells/mL) in identical circular wells (3.26 mm diameter). Higher cell densities drove greater matrix compaction, with 4-post configurations showing enhanced compaction compared to 8-post at densities ≥10^6^ cells/mL.. Scale bars: 1 mm. **(b)** Quantification of tissue compaction based on 2D projected area measurements. Data shown as mean ± SEM (*n* = 8 tissues per condition from a single batch representative of 3 independent experiments). At low densities, minimal compaction occurred in both geometries. Progressive increases in cell density enhanced compaction, with 4-post tissues showing significantly smaller final areas than 8-post tissues at densities ≥10^6^ cells/mL (\*\*\**p* < 0.001, \*\*\*\**p* < 0.0001, two-way ANOVA with Holm-Sidak post-hoc test). The increased compaction in 4-post devices at higher densities supports model predictions that anisotropic boundary conditions enhance cell contractility.

We next used the models to predict the mechanical stress field generated within tissue in each design. The directionality of mechanical stress within these tissues varies significantly based on post geometry. In the 8-post configuration, the central region of the tissue experiences an isotropic stress field, where principal stresses are equal in all directions. However, the peripheral regions display strong circumferential anisotropy, with tension aligning tangentially around the boundary. In contrast, the 4-post configuration generates an anisotropic stress field, especially in the central region, where tension is strongly aligned horizontally between opposing posts. Only small zones between neighboring posts on each side experience partial isotropy (**Fig. 3a**).

We next analyzed the magnitude of stress generated within the tissue using conventional models, including pre-stressed/pre-strained formulations and conventional mechanical models commonly implemented in commercial finite element software. We plotted the results along a horizontal transect (line A–B) spanning the tissue center between opposing posts. These classical models predicted that tension is highest in the central region of the 8-post configuration and lowest at the center of the 4-post configuration (**Fig. 3b**). Given that fibroblast activation is known to increase with tension, these classical models predict that cells in the center of an 8-post tissue should be more activated than cells in the center of a 4-post tissue.

However, our model predicted the exact opposite pattern of fibroblast activation. In the 4-post tissues, the central region exhibited the highest levels of predicted fibroblast activation, whereas in the 8-post configuration, activation was lowest in the central region (**Fig. 3c**). This reversal arises because our model accounts not only for stress magnitude but also for stress anisotropy as a regulator of cellular activation. In our model, the average of cellular contractility 𝜌_𝑘𝑘_ increases through two feedback mechanisms: one driven by the mean stress magnitude 𝜎_𝑘𝑘_, and the other by the anisotropy of the local stress field (𝜎_a_):

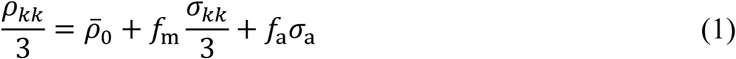

where 𝜌̅_0_ is the baseline cell contractility (in the absence of tension), and 𝑓_m_, and 𝑓_a_ are model parameters regulating the increase in cell contractility in response to the magnitude and anisotropy of the stress field, respectively.^19^ The anisotropy of the stress field, 𝜎_a_, is defined as the differences in principal stress values of the stress tensor.

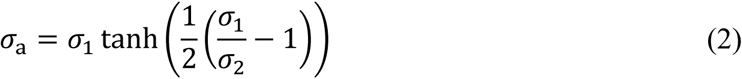

where 𝜎_1_ and 𝜎_2_ are the first and second principal components of the stress tensor. This formulation predicts cell activation and the promotion of cell contractility in regions where the stress field is anisotropic. As a consequence of this anisotropy-driven contractility, cells in the center of 4-post tissues become more activated, generating increased local force. This force, in turn, amplifies tissue-level tension in the same region (**Fig. 3d**), opposite to the stress gradient predicted by classical models (**Fig. 3b**).

Together, our simulations predict that fibroblast activation is regulated not solely by the magnitude of tensile stress, but more critically by the anisotropy of the stress field. In the next sections, we experimentally test these model predictions in an experimental model of wound compaction, in which cell density is high and significant global matrix remodeling occurs.

### Anisotropic boundaries enhance tissue compaction at high cell density

We began by measuring changes in tissue morphology and compaction across a range of cell densities. After 24-hour incubation in microwells, the compacted area of 3D fibroblast-collagen tissues in different cell densities was recorded (**Fig. 4a,b**). Projected area across different densities showed that the tissues underwent stepwise compaction responding to the boundaries created by the posts. Importantly, at this early timepoint, cells seeded at the lowest density (2.5x10^5^ cells/mL) had not yet begun to appreciably compact the collagen matrix, whereas cells at higher densities already exhibited significant matrix compaction, with the overall degree of compaction being higher as the volumetric densities of cells increased (**Fig. 4b, Fig. S1**). Above an initial cell seeding density of 10^6^ cells/mL collagen matrix, we observed that 4-post based tissues occupied a lower area than 8-post based tissues. Importantly, even when we normalized the available area of a polygon connecting the posts, cell-generated compaction in 4-post tissues yielded more inward compaction after 4 days of culture (**Fig. S2**). Side-views of OCT 3D reconstructions confirmed that for both 4-post and 8-post designs, tissues successfully detached from the surface of microwells, validating our computational model’s assumption of free compaction in all 3 dimensions (**Fig. S3**).

### Stress anisotropy drives a temporal cascade of mechanosensing from individual cell elongation to collective tissue alignment

To understand how fibroblasts integrate mechanical anisotropy across scales, we examined whether tissue-level compaction is prerequisite for cellular mechanosensing. Surprisingly, even at the lowest cell density tested (2.5 × 10^5^ cells/mL), where bulk matrix compaction was negligible and tissues failed to span between posts, individual fibroblasts exhibited significant elongation in response to boundary-imposed stress fields (**Fig. 5a-c**). This elongation was directionally biased: cells in the center of 4-post tissues showed 42% greater aspect ratio compared to those in 8-post centers (3.8 ± 0.4 vs. 2.7 ± 0.3, *p* < 0.001), demonstrating that individual cells can detect and respond to mechanical anisotropy before any appreciable tissue reorganization occurs. Nuclear elongation followed a similar trend. However, while nuclear elongation increased in fibroblasts in the center of 4-post versus 8-post tissues (**Fig. 5d,e**), differences were less stark in nuclear than in cell elongation.

**Figure 5.**
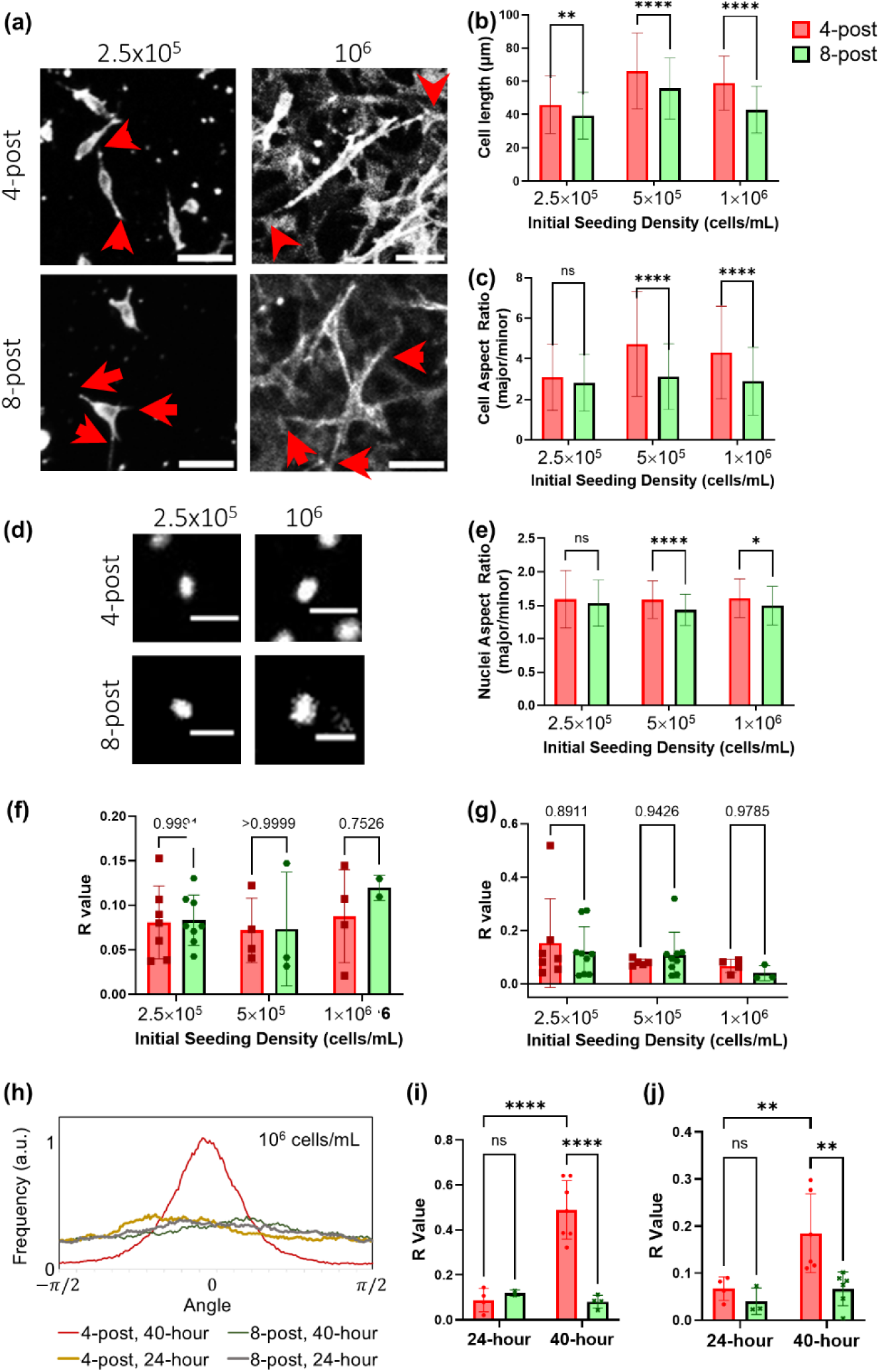
Temporal hierarchy of mechanosensing reveals individual cell elongation precedes collective alignment in response to stress anisotropy. **(a-c)** Representative cell morphologies at 24 hours across initial cell densities. **(a)** Representative immunofluorescence images showing F-actin in the central region of 4-post and 8-post tissues. Arrows: cell protrusions. Scale bar: 50 μm. **(b)** Quantification of cell length showing enhanced elongation in 4-post tissues at all densities above 0.25 × 10^6^ cells/mL. **(c)** Cell aspect ratio analysis confirms greater elongation in anisotropic (4-post) versus isotropic (8-post) stress fields. **(d-e)** Nuclear morphology mirrors cellular elongation patterns at 24 hours. **(d)** Representative DAPI-stained nuclei. Scale bar: 20 μm. **(e)** Nuclear aspect ratio quantification across cell densities, demonstrating geometry-dependent differences. **(f-g)** Collective alignment lags behind individual elongation. Despite significant differences in cell elongation at 24 hours, alignment (R-value) shows no difference between geometries across all densities tested. R-value ranges from 0 (random orientation) to 1 (perfect alignment). **(h-j)** Extended culture reveals emergence of collective organization. **(h)** Representative images at 40 hours (10^6^ cells/mL) showing aligned cells in 4-post tissues versus mixed orientations in 8-post tissues. **(i)** Temporal evolution of cellular R-value demonstrates progressive alignment in 4-post tissues between 24-40 hours. **(j)** Nuclear alignment follows similar temporal pattern, confirming coordinated tissue-level reorganization. Data presented as mean ± SEM (*n* ≥4 tissues per condition from 3 independent experiments). Statistical significance: *p < 0.05, **p < 0.01, ****p < 0.0001 by two-way ANOVA with Holm-Sidak post-hoc test for pairwise comparisons.

The relationship between cell density and mechanosensing revealed distinct scaling behaviors for different cellular processes. Individual cell elongation increased modestly from the lowest density (2.5 × 10^5^ cells/mL) to intermediate densities (5 × 10^5^ cells/mL) but then plateaued: cells at 10^6^ cells/mL exhibited similar elongation to those at half this density (**Fig. 5a-c**). In contrast, tissue compaction scaled continuously with cell density, showing progressive increases at each density increment without saturation (**Fig. 4**). This indicates that individual cellular mechanosensing operates through mechanisms distinct from, and more sensitive than, bulk tissue mechanics.

We observed a temporal hierarchy in how cells translate mechanical anisotropy into organized tissue architecture. At 24 hours, despite pronounced differences in individual cell elongation between 4-post and 8-post geometries, both configurations showed similarly low collective alignment. Collective alignment was quantified through the R-value (**Fig. S4**), which quantifies the frequency at which cell orientation follows a given angle (**Fig. S4**). Quantitative analysis at the 24 hour timepoint was consistent with the qualitative observation that fibroblast alignment was largely random (R-values: <0.1; **Fig. 5f,g**). However, extending culture to 40 hours revealed dramatic reorganization specifically in anisotropic environments: 4-post tissues developed strong uniaxial alignment (R-value: 0.49 ± 0.12) while 8-post centers remained largely isotropic (R-value: 0.081 ± 0.029; *p* < 0.01; **Fig. 5h-j**).

This temporal cascade from individual elongation (≤24 hours) to collective alignment (24-40 hours) reveals that mechanosensing operates through sequential, scale-dependent processes. Initially, individual cells detect stress anisotropy through local matrix interactions, elongating along principal stress directions. This primary response requires minimal cell-cell coordination and occurs even at low densities where cells are largely isolated. Subsequently, elongated cells undergo rotational reorientation, aligning their long axes to establish tissue-level organization. This secondary process involves progressive cell-cell mechanical communication and matrix remodeling, creating positive feedback that amplifies initial anisotropic cues.

This mechanosensing hierarchy persisted even at high cell densities (10^6^ cells/mL) where extensive cell-cell contacts might be expected to dominate over matrix-mediated signals. The preservation of geometry-dependent responses across a 4-fold range in cell density (2.5 × 10^5^ to 10^6^ cells/mL) demonstrates that stress anisotropy sensing represents a fundamental cellular capability that operates robustly across diverse tissue contexts. Higher cell densities accelerated but did not bypass this temporal sequence, suggesting that the progression from individual to collective mechanosensing reflects intrinsic constraints on how cells process and integrate mechanical information across scales.

### Stress anisotropy culminates in enhanced myofibroblast differentiation following the morphological cascade

Having established that stress anisotropy drives sequential morphological changes, from individual cell elongation to collective alignment, we investigated whether this mechanical sensing cascade ultimately governs fibroblast phenotypic fate. The fibroblast-to-myofibroblast transition (FMT) represents a critical phenotypic switch characterized by upregulation of α-smooth muscle actin (α-SMA) and enhanced F-actin stress fiber formation.^29–32^

Consistent with our observed temporal hierarchy, FMT markers showed delayed kinetics relative to morphological changes. At 24 hours, when cells had already elongated significantly in response to stress anisotropy, we detected no difference in α-SMA expression between 4-post and 8-post tissue centers across multiple cell densities (**Fig. S5**). This temporal mismatch suggested that while cells rapidly sense and morphologically respond to mechanical anisotropy, the transcriptional and translational machinery driving phenotypic transition requires additional time or sustained mechanical stimulation to engage.

To capture the full progression of mechanically-induced FMT, we extended our analysis to 96 hours using high-density tissues (2×10^7^ cells/mL) where both morphological reorganization and mechanical feedback would be maximized. This extended culture revealed striking geometry-dependent phenotypic differences that precisely matched our computational predictions. In the central regions of 4-post tissues, where our model predicts maximum stress anisotropy, we observed ∼40%higher α-SMA intensity compared to 8-post centers, where stress is nearly isotropic (4-post: 101 ± 25 A.U. vs. 8-post: 74 ± 13 A.U., *p* < 0.05; **Fig. 6a,f**). F-actin stress fibers showed similar enhancement, withgreater intensity in anisotropic versus isotropic regions (4-post: 76 ± 16 A.U. vs. 8-post: 56 ± 15 A.U., *p* < 0.05).

**Figure 6.**
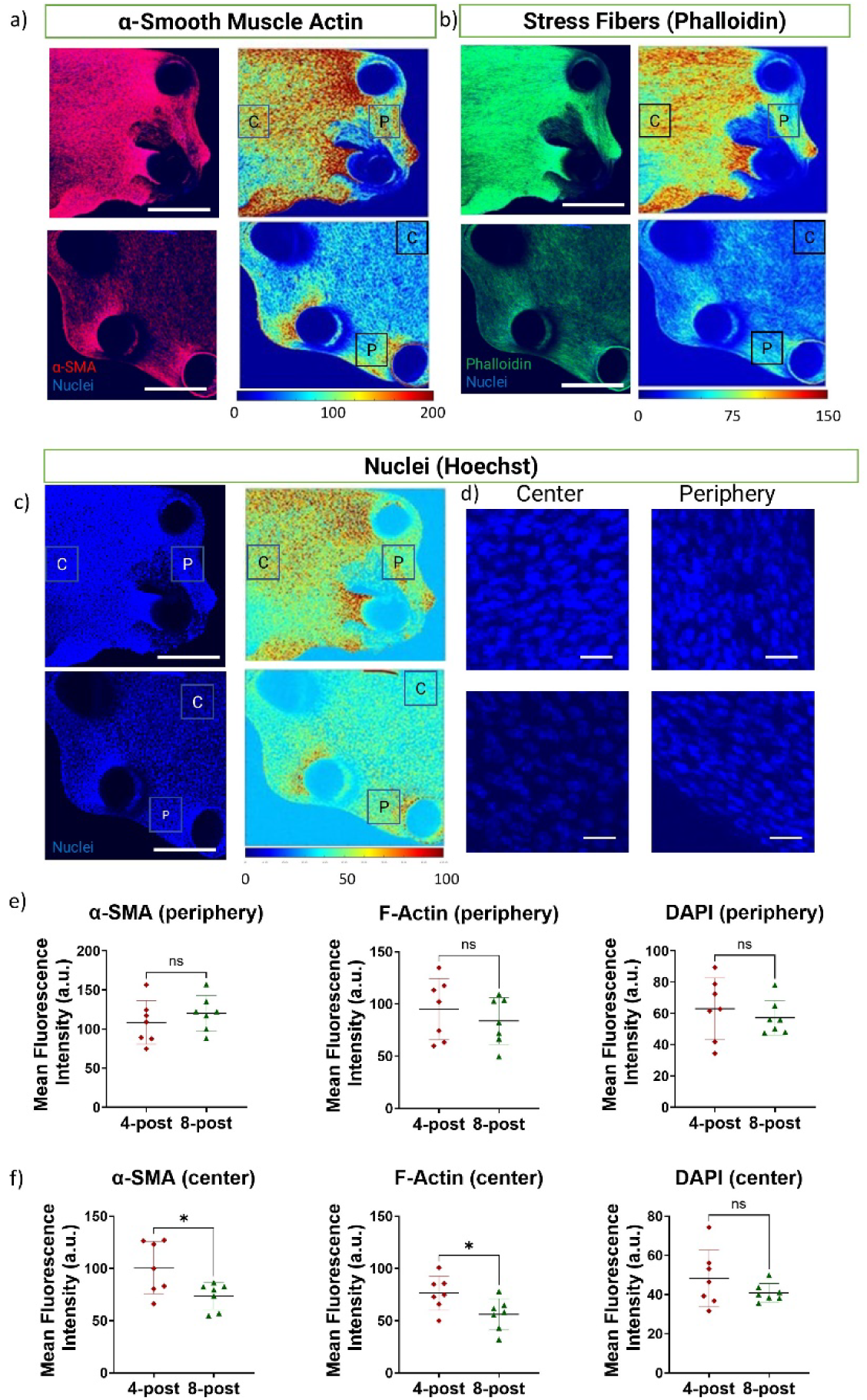
Stress anisotropy drives spatially patterned myofibroblast differentiation consistent with computational predictions. **(a-c)** Representative maximum intensity projections of whole tissues after 96 hours of culture (initial density: 2×10^7^ cells/mL) showing spatial distribution of **(a)** α-smooth muscle actin (α-SMA, red), **(b)** F-actin (green), and **(c)** nuclei (DAPI, blue). Heat map overlays indicate relative fluorescence intensity. Note enhanced α-SMA and F-actin expression in the center of 4-post tissues and peripheral regions of both geometries, corresponding to regions of high stress anisotropy predicted by the computational model (cf. Fig. 3a). **(d)** High-magnification views of nuclear organization in central (“C”) and peripheral (“P”) regions of interest (ROIs), revealing comparable cell densities but distinct organizational patterns between geometries. **(e)** Quantification of fluorescence intensity in peripheral ROIs shows equivalent α-SMA and F-actin expression between 4-post and 8-post tissues (n.s., not significant), consistent with similar stress anisotropy at tissue boundaries in both geometries. Nuclear density (DAPI) confirms comparable cell numbers. **(f)** Quantification in central ROIs reveals significantly higher α-SMA and F-actin expression in 4-post versus 8-post tissues (****p < 0.05), despite similar nuclear densities, demonstrating that stress anisotropy rather than cell density drives myofibroblast differentiation. Data represent mean ± SEM from *n* = 8 tissues per condition across 3 independent experiments. Statistical analysis by unpaired two-tailed t-test. Images acquired as 100 z-slices at 5.09 μm intervals and displayed as maximum intensity projections. Scale bars: (a-c) 500 μm; (d) 50 μm.

Spatial analysis provided further validation of our anisotropy-driven activation model. In peripheral regions of both 4-post and 8-post tissues, where the model predicts comparable stress anisotropy due to circumferential tension around the tissue boundary, we observed statistically indistinguishable levels of both α-SMA and F-actin expression (**Fig. 6a,e**). This spatial pattern— high activation in both the 4-post center and all peripheral regions, but low activation in the 8-post center—is unlikely to be explained by differences in cell density, as overall nuclear intensity remained comparable between central regions (4-post: 48 ± 15 A.U. vs. 8-post: 40 ± 5 A.U., *p* = 0.23; **Fig. 6c,f**), nor by classical models based solely on stress magnitude.

The 72-hour delay between initial mechanical sensing (cell elongation at 24 hours) and full phenotypic commitment (α-SMA upregulation at 96 hours) reveals that FMT represents the culmination of a multi-stage mechanotransduction cascade. This progression, from rapid morphological response to gradual collective reorganization to eventual phenotypic transition, demonstrates how cells integrate mechanical signals across multiple timescales to make fate decisions. The requirement for sustained anisotropic stress to drive FMT has important implications for understanding both normal wound healing, where transient mechanical cues promote appropriate repair, and pathological fibrosis, where persistent mechanical anisotropy may lock cells in an activated state.

These results establish stress anisotropy, rather than stress magnitude alone, as the primary mechanical driver of fibroblast activation in 3D tissues. Even in dense tissues where cell-cell interactions are extensive and matrix remodeling is pronounced, the directionality of mechanical stress continues to govern cell fate, highlighting the fundamental importance of tissue geometry in regulating fibroblast biology.

## 3. Discussion

Our results reveal that fibroblast mechanosensing of stress anisotropy operates through a hierarchical temporal cascade, with individual cell elongation preceding collective alignment, which in turn precedes myofibroblast differentiation. This temporal progression, from hours to days, suggests integration of mechanical information across multiple timescales as cells make phenotypic decisions. Anisotropy sensing persists even in high-density tissues where extensive cell-cell contacts and matrix remodeling might be expected to mask mechanical signals from tissue boundaries. These findings establish stress directionality, rather than magnitude alone, as a fundamental regulator of fibroblast fate in 3D tissues.

### Temporal hierarchy reveals multiscale mechanosensing mechanisms

The temporal hierarchy identified begins with individual cell elongation (≤24 hours), and progresses to collective alignment (24-40 hours) and phenotypic transition (72-96 hours) (**Fig. 7**). This suggests that mechanotransduction operates through distinct, sequential processes and is thus consistent with emerging evidence that cells employ multiple mechanosensing mechanisms operating at different timescales.^22,34^ The rapid initial elongation likely reflects immediate cytoskeletal reorganization in response to local matrix forces, potentially mediated by stretch-activated channels or integrin clustering. The subsequent collective alignment phase requires cell-cell mechanical communication and coordinated matrix remodeling,^27,35–38^ creating the positive feedback loops we previously described between cellular contractility and ECM organization.^22^ Finally, the delayed myofibroblast transition suggests involvement of transcriptional programs that require sustained mechanical stimulation, consistent with the known kinetics of mechanically-induced gene expression.^39–42^

**Figure 7.**
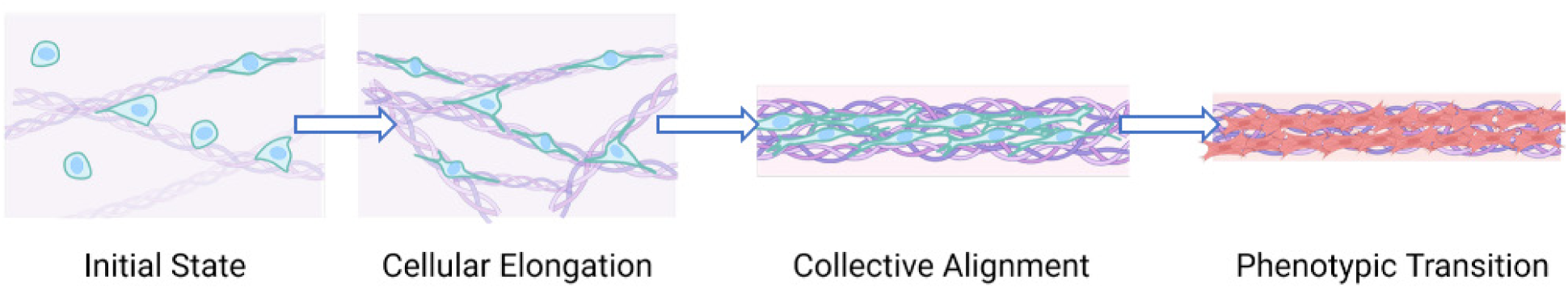
Temporal cascade of mechanosensing reveals how stress anisotropy drives progressive fibroblast activation in 3D tissues. Schematic timeline illustrating the hierarchical sequence of cellular responses to anisotropic (4-post) versus isotropic (8-post) mechanical boundary conditions. Three distinct phases emerge: **(i) Early phase (0-24 h)**: Individual cells detect stress anisotropy and elongate along principal stress directions, independent of bulk tissue compaction or cell density. **(ii) Intermediate phase (24-40 h)**: Elongated cells undergo collective reorientation to establish tissue-level alignment, requiring cell-cell mechanical communication and matrix reorganization. **(iii) Late phase (40-96 h)**: Sustained exposure to anisotropic stress culminates in myofibroblast differentiation marked by α-SMA upregulation and enhanced actin polymerization. In isotropic conditions, this progression is suppressed. Higher cell densities accelerate but do not bypass this temporal sequence, suggesting an intrinsic program whereby cells must integrate mechanical signals across multiple timescales before committing to phenotypic changes.

Our observation that cells sense anisotropy even at densities where bulk tissue compaction is minimal (2.5 × 10^5^ cells/mL) indicates that individual cells can detect far-field mechanical boundaries through the collagen network. This finding supports theoretical predictions that fibrous ECM networks can transmit mechanical information over distances exceeding typical cell-cell spacing.^26,33,43^ The persistence of anisotropy sensing at high densities (10^6^ cells/mL), where cell-cell mechanical networks become prominent, demonstrates that boundary-imposed stress fields continue to influence cell behavior even when local cell-generated forces might be expected to dominate.

### Reconciling stress magnitude and anisotropy in fibroblast activation

Our anisotropy-sensing model correctly predicted fibroblast activation patterns opposite to those expected from classical mechanics. While conventional models predict highest stress and activation in 8-post centers due to greater mechanical resistance, we observed maximum activation in 4-post centers where stress is most anisotropic. This apparent paradox is resolved by recognizing that cells in anisotropic stress fields can generate higher contractile forces through polarized cytoskeletal organization, ultimately producing higher local stress despite lower boundary resistance. This mechanism, captured by our model’s dual feedback terms (Eq. 1), demonstrates how cellular mechanosensing of directionality can override effects of stress magnitude.

The success of our model in predicting both spatial patterns (center vs. periphery) and phenotypic outcomes (α-SMA expression) supports its fundamental hypothesis that fibroblasts integrate both stress magnitude and anisotropy to regulate their contractility. The model’s ability to capture these behaviors using relatively simple phenomenological relationships suggests that anisotropy sensing may be a fundamental property of the actomyosin cytoskeleton, potentially arising from the inherent tendency of stress fibers and microtubules to align with principal stress directions.^18,22,28^

### Implications for wound healing and fibrosis

The temporal cascade we identified has important implications for understanding wound healing dynamics. During normal repair, the sequential nature of mechanosensing may serve as a regulatory checkpoint, allowing tissues to distinguish between transient mechanical perturbations and sustained injury requiring myofibroblast activation. The 72-96 hour delay before full α-SMA upregulation provides a window for resolution of mechanical stress through tissue reorganization alone, without committing to the more dramatic phenotypic changes associated with myofibroblast differentiation.

In pathological fibrosis, however, persistent mechanical anisotropy arising from non-resolving inflammation, continued tissue damage, or architectural disruption could lock cells in a state of sustained activation.^4,15–17^ Our results suggest that the geometry of fibrotic lesions, not just their stiffness, may critically influence disease progression. Therapies targeting the mechanical environment might therefore benefit from considering stress directionality alongside the more commonly studied parameter of matrix stiffness.^12–14^

### Design principles for tissue engineering

Our findings highlight the importance of geometric design in tissue engineering applications. The dramatic differences in cell behavior between 4-post and 8-post configurations (despite identical materials, cell sources, and culture conditions) demonstrate that boundary geometry alone can determine tissue phenotype. This has immediate implications for engineering tissues where controlled fibroblast activation is desired (e.g., wound healing scaffolds) or must be avoided (e.g., anti-fibrotic implants).^44,45^

For engineered cardiac or skeletal muscle, where some degree of ECM organization is beneficial, but excessive fibroblast activation impairs function, our results suggest that moderately anisotropic geometries might optimize the balance between tissue organization and myofibroblast suppression. The HASTE fabrication platform we developed enables rapid iteration of such geometric designs, facilitating systematic optimization of boundary conditions for specific applications.

### Mechanistic insights and future directions

While our study establishes the phenomenology of anisotropic mechanosensing, the molecular mechanisms remain to be fully elucidated. Several non-exclusive mechanisms could contribute to anisotropy detection: (1) differential activation of mechanosensitive ion channels under polarized versus isotropic stress;^46^ (2) anisotropic nuclear deformation affecting chromatin accessibility and gene expression;^28^ (3) polarized integrin clustering and focal adhesion maturation;^47^ or (4) microtubule-actomyosin crosstalk that we previously showed regulates directional contractility.^19,22^ Future studies employing targeted molecular perturbations will be needed to dissect these mechanisms.

The relationship between our findings in fibroblasts and mechanosensing in other cell types also warrants investigation. Recent studies in stem cell-derived cardiomyocytes suggest that prestress influences maturation,^20,48^ but whether these effects arise from stress magnitude or anisotropy remains unclear. Our platform provides a tool to systematically address this question across cell types.

### Study limitations and technical considerations

Several limitations should be considered when interpreting our results. First, the inability to track individual cells in dense 3D cultures prevented us from distinguishing cell-autonomous α-SMA upregulation from population heterogeneity. Second, while we focused on serum-containing conditions to maintain cell viability over 96 hours, factors within serum, including TGF-β, may influence the kinetics of myofibroblast differentiation. Future studies in defined media could isolate purely mechanical effects. Third, our model treats tissues as continua, potentially missing discrete cellular behaviors that emerge from stochastic cell-level decisions.

Despite these limitations, the robustness of our findings across a 40-fold range in cell density, the agreement between experiments and computational predictions, and the clear temporal progression of mechanosensing events provide strong evidence that stress anisotropy fundamentally regulates fibroblast biology in 3D tissues.

## 4. Conclusions

Our work establishes that fibroblasts sense and respond to stress anisotropy through a hierarchical spatiotemporal cascade that culminates in myofibroblast differentiation. The sequence, from rapid individual cell elongation to collective tissue organization to phenotypic commitment, reveals how cells integrate mechanical information across timescales to make fate decisions. Our findings demonstrate that tissue geometry, through its effect on stress directionality, can be as important as biochemical factors in controlling fibroblast activation. These results provide insight into wound healing and fibrosis, and offer design principles for engineering tissues with controlled fibroblast phenotypes.

## 5. Methods

### Device design and fabrication

#### 3D Printing of Master Molds

Micropost array devices were designed in SolidWorks 2020 (Dassault Systèmes, Waltham, MA). Each platform contained 192 individual devices arranged in a 12×16 array. Individual devices consisted of circular wells (3.26 mm diameter, 1.25 mm depth) containing either 4 or 8 microposts. Posts were designed with 200 μm shaft diameter and 1.125 mm height, topped with overhanging caps (300 μm diameter, 125 μm thickness). Inter-post spacing was 1.86 mm between opposing posts and 1.31 mm between adjacent posts. Master molds were printed using a Form 3 3D printer (Formlabs, Somerville, MA) with Clear Resin (Formlabs, Somerville, MA) at 25 μm layer resolution. Post-processing involved dual 15-minute washes in isopropyl alcohol, air drying, and 30-minute UV curing, followed by careful removal of support structures.

#### HASTE Fabrication Protocol

PDMS devices were fabricated using our modified Hydrogel-Assisted Stereolithographic Elastomer (HASTE) protocol. ^20^ Briefly, agar molds (1.5% w/v in tap water, Fisher Scientific) were prepared by dissolving agar at 100°C, cooling to 60°C, and adding Triton X-100 to 0.2% final concentration. In comparison cases, no Triton X-100 was added. This surfactant-modified agar was poured over 3D-printed masters and allowed to gel at room temperature (10 min), followed by 4°C (20 min). After demolding, the agar-negative molds were used for PDMS casting.

#### Compliant Micropost Formation

Two-step PDMS casting was employed to create compliant posts within rigid wells. First, a soft PDMS mixture (elastic modulus ∼130 kPa) was prepared by combining Sylgard 184 (1:10 base:crosslinker ratio, Dow Corning) with Sylgard 527 (1:1 Part A:Part B ratio) at a 1:4 ratio.^21,53^ This mixture was degassed, poured into agar molds, and partially cured at 37°C for 2 hours. Subsequently, standard Sylgard 184 (elastic modulus ∼2 MPa) was added to form rigid wells and backing, with final curing at 60°C overnight. Devices were removed from agar molds and underwent terminal curing at 60°C for an additional 24 hours.

### Cell culture

NIH/3T3 fibroblasts (gift from Spencer Lake, Washington University in St Louis) were cultured in Dulbecco’s modified Eagle’s medium (DMEM; Gibco, USA) supplemented with 10% Fetal Bovine Serum (Midsci, USA), 1% Non-essential amino acid (NEAA; Gibco, USA), 100U/mL Penicillin-streptomycin (Gibco, USA), and 1% GlutaMAX (Gibco, USA) at 37°C and 5% CO2. Cells were passaged every 3 days (using 0.05% trypsin/EDTA) at a 1:10 ratio. To form micro-tissues, precise counts of viable fibroblasts were obtained using Trypan blue exclusion on an automatic cell counting machine (Invitrogen Countess 2).

### 3D microtissue formation

Four different cell densities were used: 2.5x10^5^, 5x10^5^, 10^6^, and 2x10^7^ cells/mL. Cells were mixed on ice with rat-tail collagen (Gibco A1048301) with final concentration 1.5 mg/ml, 0.5 mM Hepes (Gibco) in DMEM, and neutralized with NaOH (final concentration of 12.5 mM). The seeding to devices was carried out on top of an icebox and then centrifuged at 300 RCF for 3 minutes in a chilled centrifuge (4°C). Following 15 minutes incubation at 37°C to allow collagen to crosslink, media (DMEM with10% FBS) was added. The media was replenished every 2 days. Bright field images were taken using Nikon Eclipse TsR at 3 hours, 1 day, and 4 days after seeding the engineered fibroblast tissues.

### 3D morphology analysis by optical coherence tomography

A customized Spectral Domain - Optical Coherence Tomography (SD-OCT) system^54,55^ recorded the 3D morphology of the microtissues on the day of seeding as well as on day 4 before fixation. In brief, a superluminescent diode (EXALOS, EXC250023-00, central wavelength: 1300 nm, spectral range: 180 nm) provided the light source of OCT. A spectrometer (Wasatch Photonics, Cobra 1300) comprising a 2048-pixel InGaAs line-scan camera (Sensors Unlimited, GL2048) allowed a 147 kHz A-scan rate at its maximum. Each 3D-OCT image consisted of 600 A-scans per B scan and 600 B scans to cover a field of view of 2.92 mm x 2.76 mm with resolutions of ∼2.83 μm (transverse) and ∼3.95 μm (axial) in DMEM (refractive index = 1.345). The imaging process took about 31 seconds with an exposure time of 60 μs. Using ImageJ, voxel size in x-y-z dimensions was normalized isotropically.^56^ Then, using a machine-learning-based segmentation of the tissue was done to remove minor noises before volume measurement.

### Immunofluorescence microscopy of microtissues

Microtissues were fixed in 4% paraformaldehyde (PFA) in Dulbecco-phosphate buffered saline (dPBS) solution for 20 minutes at room temperature. Following fixation, the microtissues were washed 2 times with dPBS for 5 minutes each at room temperature, followed by a final wash overnight at 4°C. Selected devices with microtissues were carefully cut and transferred to an 8-well chamber slide to perform immunostaining. Samples were permeabilized with 0.25% Triton X-100 for 15 min and blocked using 5% Normal Goat Serum for 1 hour at room temperature. Rabbit recombinant monoclonal α-smooth muscle actin antibody (AB124964, Abcam) was incubated overnight at 4 °C. After washes, secondary antibody (Alexa Flour 647, A21235, Invitrogen), Phalloidin-488 (A12379, Thermo Fisher), and cell nuclei stain (Hoechst 33342, ThermoFisher) were incubated at room temperature for 2 hours. After washing steps, the samples were then kept in dPBS before being imaged within 12 hours.

Samples were imaged on an Olympus confocal microscope (Fluoview FV1200, Tokyo, Japan) using a 10X objective in XY scan mode at a sample speed of 20.0 μs/pixel. Confocal scans were performed using line sequential mode, with line Kalman integration on, with a count of 8. All z-slices were taken with an interval of 5 µm to perform analysis with maximum intensity projections.

### Cell and nucleus outline analysis

To quantify cell morphology (cell and nucleus aspect ratio, nuclei orientation, and cell length in the tissue), ImageJ was used. The following steps were followed to analyze binarized images of individual cells. Firstly, region of interest with size of 1 mm x 1 mm at the center of tissue was cropped from maximum Z-projection of each tissue specimen. Next, each cell was manually traced using the Freehand Selection tool to outline the boundary of the cell. Only fully visible and isolated cells were selected to avoid segmentation artifacts from neighboring cell features. Double-checking with nuclei position helped to distinguish the isolated cell boundary. After selecting the cell outline, the cell morphology was measured using the “Fit ellipse” function under the Analyze > Set Measurements menu, ensuring the options for “Fit ellipse” and “Shape descriptors” were enabled. Then, based on the fitted ellipse, the major axis and minor axis lengths were recorded. The cell length was defined as the major axis length that has been adjusted to micrometer scale. The aspect ratio (AR) was calculated as the ratio of major axis to minor axis (AR = Major/Minor), to provide a measure of cell morphology. The angle of fitted ellipse from nuclei is then calculated for the R value of orientation distribution, which refers to the resultant vector length. This value quantifies the degree of orientation coherence among the population.

### Fiber orientation distribution analysis

To see whether the cells have responded to mechanical anisotropy in collective way, we investigated the overall distribution of individual cells and nuclear alignment. We quantified the fibroblast alignment in the center of the cultured engineered fibroblast tissue by extracting maximum intensity projection of a certain number of confocal layers of the visualized alpha-SMA. The in-plane orientation from the center area, 1 mm x 1 mm (**Fig. S6**), was analyzed using the OrientationJ plug-in for ImageJ (available from the Biomedical Imaging Group, Switzerland; http://bigwww.epfl.ch/demo/orientation/). The measurement was performed using a Gaussian-weighted gradient with a local window σ of 2 pixels. The orientation distribution was then shown as a histogram of orientation value, -90°to +90°, versus occurrence value without additional binning from the plugins output (interval: 1°).

To ensure positive interpretation, we use conventional formulation approach to compute R based on directional concentration of orientation distributions. The R value represents the resultant vector length derived from weighted sine and cosine projections. Intensity values were employed as weights for calculations across the full orientation spectrum of -90° to +90°.

### **α**-smooth muscle actin and f-actin expression analysis

Custom MATLAB code (supporting file) was written to analyze the mean fluorescent intensity values of the different fluorescent markers within different points of the tissues. Briefly, each of the wavelengths was saved into separate TIFFs using the built-in mean intensity function in Olympus Life Science’s FV10-ASW V4.2 Viewer. Using the custom MATLAB code, the three TIFFs corresponding to the three wavelengths of a single location on the tissue were imported into MATLAB and visualized for the user. The user then defined an N-by-N square ROI on the location of the image to be analyzed. The mean intensity values within the ROI for the three wavelengths were then calculated and compiled into an Excel spreadsheet for quantitative analysis.

### Computational Model

The computational simulations were based on our recently calibrated model,^19^ which predicts how fibroblasts regulate actomyosin-based activation in response to mechanical cues from the extracellular environment. The full details of the model are provided in the Supplementary Information (SI) of our recent work.^19,27^ The model treats fibroblasts as an active contractile force-generating element whose contractility increases in response to mechanical tension via two feedback mechanisms (Eq. (1)): (i) a tension magnitude-dependent pathway, in which the mean contractility 𝜌_𝑘𝑘_⁄3 increases with mean tension 𝜎_𝑘𝑘_⁄3, and (ii) a tension anisotropy-dependent pathway, in which mean contractility is increased in regions where tension is directionally biased (anisotropic). As a result of this feedback mechanism, cell contractility increases with both the magnitude and anisotropy of tension.

The model was implemented within a three-dimensional (3D) finite element framework using a custom UMAT (user material subroutine) in Abaqus. To simulate tissue contraction, circular tissue geometries were generated, with the initial configuration shown in Fig. 3A. Due to geometric symmetry, only one quarter of the circular domain was modeled. Micropillars were modeled as cylindrical structures with a cap at one end. The opposite end of each pillar was fully fixed to mimic the experimental setup. Contact interactions between the outer surface of each pillar and the tissue were defined using surface-to-surface contact constraints in our finite element simulations.

For the four-pillar configuration, the quarter-domain tissue mesh contained 36,272 linear hexahedral elements of type C3D8, resulting in a full-tissue mesh of 145,088 elements. For the eight-pillar configuration, the quarter-domain tissue mesh consisted of 9,630 C3D8 elements (38,520 in the full geometry). In both cases, each micropillar was meshed with 2,170 hybrid linear hexahedral elements of type C3D8H. Mesh densities in both configurations were selected based on a mesh convergence study to ensure numerical accuracy and solution convergence.

Contraction of the tissues resulted in distinct final shapes in the eight-pillar and four-pillar configurations, as shown in **Fig. 3**. For each case, we computed the spatial distribution of cell contractility 𝜌, tension 𝜎, and tension anisotropy 𝜎_a_ (**Fig. 3**).

### Statistical analysis

We used GraphPad Prism 8.3.1 for statistical analysis. To compare two groups, unpaired t-test was used to quantify two-tailed P-value. For multiple groups, ordinary one-way ANOVA was used before applying post-hoc pair-wise Holm-Sidak mean comparison test. Differences with *p* < 0.05 were considered statistically significant. * = p < 0.05; ** = p < 0.005; *** = p < 0.0005; **** = p < 0.0001.

## 6. Acknowledgements

We thank Dr Ruth Okamoto and the staff of the Jubel Makerspace at Washington University in St Louis for assistance and guidance on 3D printing. We thank Dr Barbara Semar and Dr Michael Vahey (Washington University in Saint Louis) for assistance and guidance with confocal microscopy. We thank Dr Delaram Shakiba (Johns Hopkins University) and Dr. Jose Almeida (Washington University in Saint Louis) for help with troubleshooting staining and imaging. F. A. and G. M. G. acknowledge the support by the National Institute Of Arthritis And Musculoskeletal And Skin Diseases of the National Institutes of Health under Award Number R01AR084243. N.H. acknowledges support from the National Heart, Lung and Blood Institute of the National Institutes of Health under Award Number RO1HL159094, and the Nationa Science Foundation under CAREER Award 2338931. We also acknowledge support from a McDonnell Academy Fellowship (G.R.). The content is solely the responsibility of the authors and does not necessarily represent the official views of the National Institutes of Health or the National Science Foundation.

