## Supplementary Information for "3D-printing-assisted, microfabricated devices reveal hierarchical and temporal mechanosensing in high-density fibroblast culture"

### Supplemental Information

#### Supplementary Figures

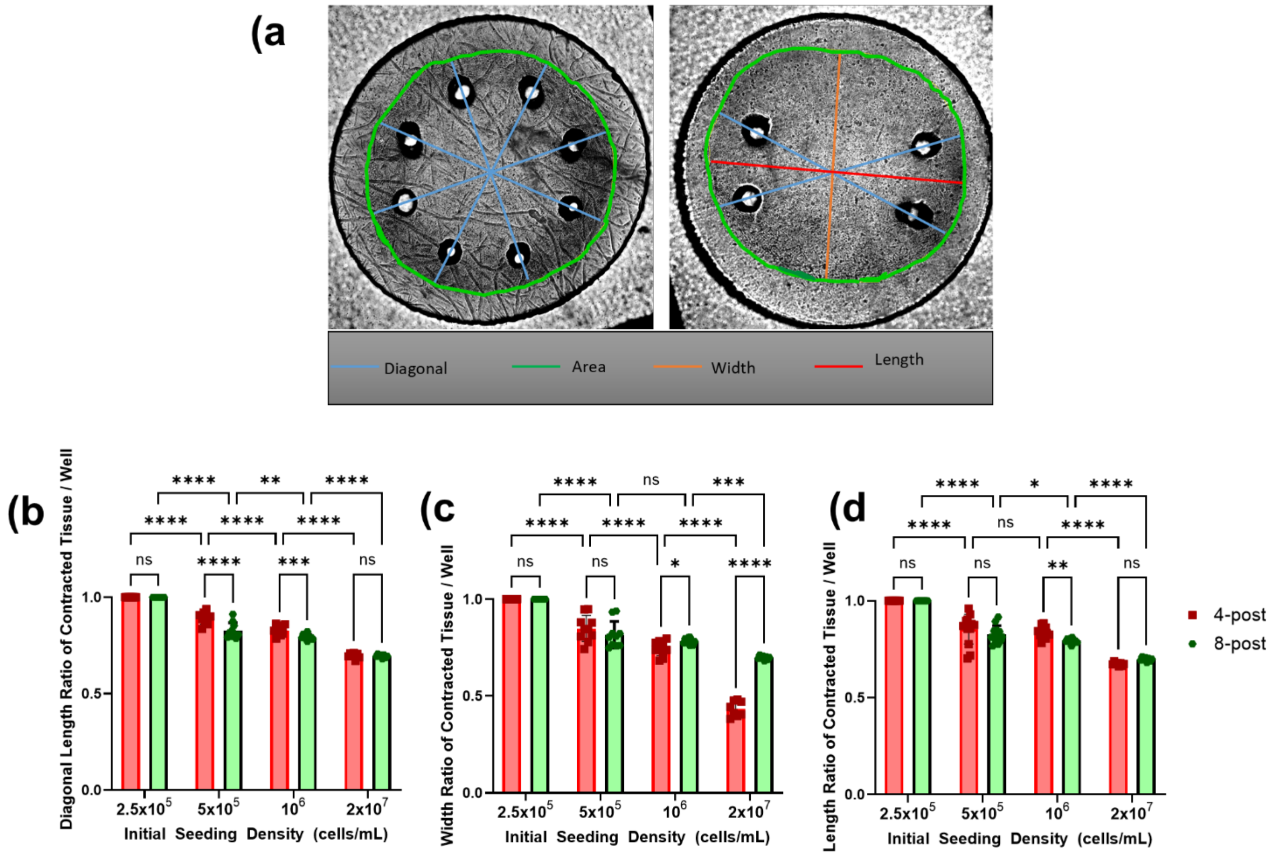


Figure S1. Contraction data was visualized by measuring ratio of each contracted tissue compared to the overall dimension of each well. a) Representative micrographs of (left) 8-post and (right) 4-post tissue with diagonal dimensions, area, width and length denoted. b-d) Quantitative analysis of b) average diagonal length, c) average tissue width, and c) average tissue length. Values for diagonal length, width and length are normalized to the diameter of the microwell containing the posts. Data are from three representative batches (total of n = 8 tissues). Data were analyzed from brightfield micrographs of engineered fibroblast tissues cultured for 24 hours. Each data point represents one tissue. Error bars: *SD*. Statistical significance: *$p < 0.05$, **$p < 0.01$, ***$p < 0.001$, ****$p < 0.0001$, ANOVA followed by Holm-Sidak post-hoc test for pairwise comparisons.

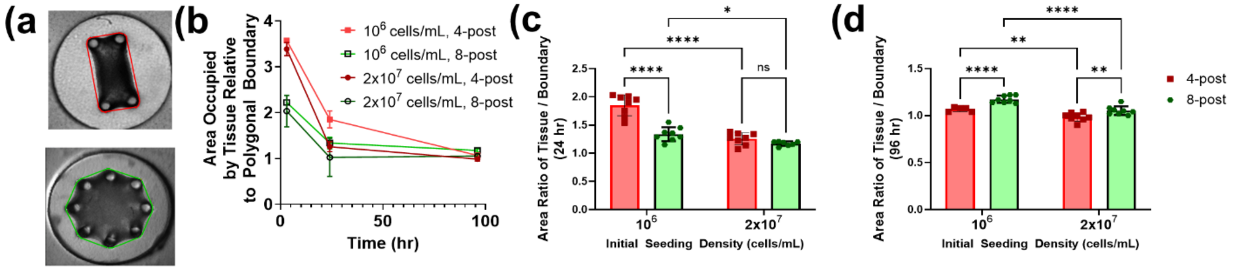


Figure S2. 2D-projected measurement of tissue-generated compaction. Because the area available within a polygon connecting the posts together (e.g., area that a tissue would occupy if it stopped compacting once it experienced the mechanical resistance of the posts), area values are normalized to the area of the polygon connecting the posts. (a) Depiction of the polygon connecting the posts. (b) Dynamics of compaction of 4 and 8-post tissues seeded with two different fibroblast densities. (c-d) comparison between 4-post and 8-post tissues for area occupied by tissue, normalized the the area of the polygon connecting posts, at c) 24 hr and d) 96 hr following cell seeding.Error bars: *SD*. Each datapoint in panel a) is the mean of $n = 8$ tissues. Statistical significance: *$p < 0.05$, **$p < 0.01$, ***$p < 0.001$, ****$p < 0.0001$, ANOVA followed by Holm-Sidak post-hoc test for pairwise comparisons.

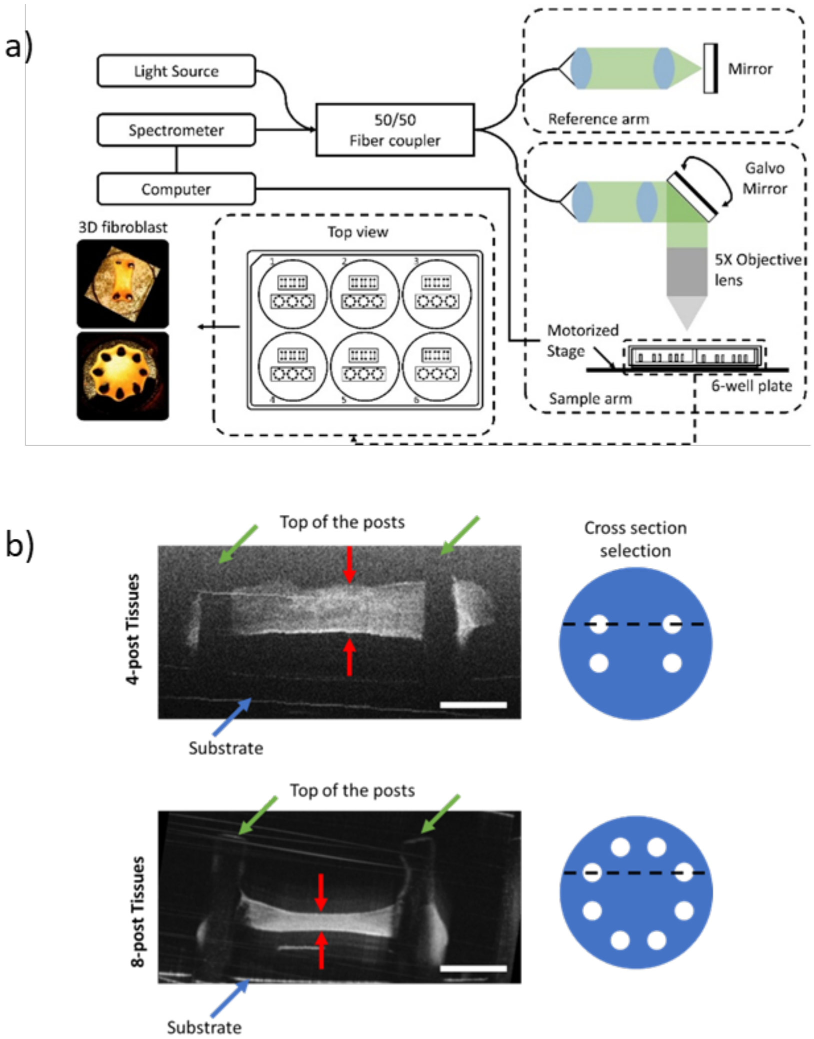

Figure S3. Optical Coherence Tomographic 3D reconstructions to assess vertical position of engineered tissue in microwells. a) Systematic of Optical Coherece Tomography (OCT) imaging system with central wavelength of 1300nm and spectral wavelength 180nm and 16.7 kHz A-scan rate (b) Side-views of 3D reconstructed OCT images of representative 4-post (top) and 8-post (bottom) microtissues. Scale Bar: 500 µm (b).

a)

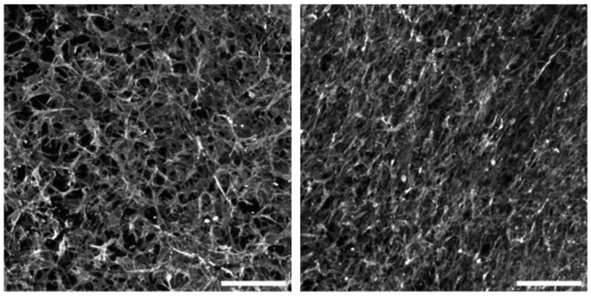

| R-value | 8-post_40-hour | 4-post_40hour |
|---|---|---|
| | 0.11 | 0.64 |

b)

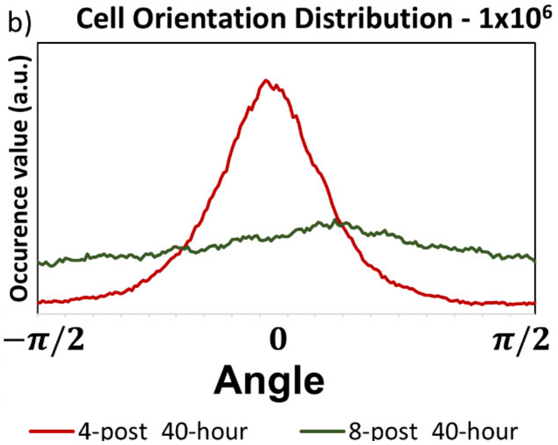


Figure S4. Representative data for orientation R values. a) Representative micrographs depicting orientation distribution of α-smooth muscle actin at the center of 8-post (left) and 4-post (right). Computed R values are depicted under the micrographs in (a). (b) Orientation distribution curves. Note that very sharp peak (4-post tissues, red) has R value closer to 1, while a fully uniform distribution (8-post tissue, green) has anR value closer to0.

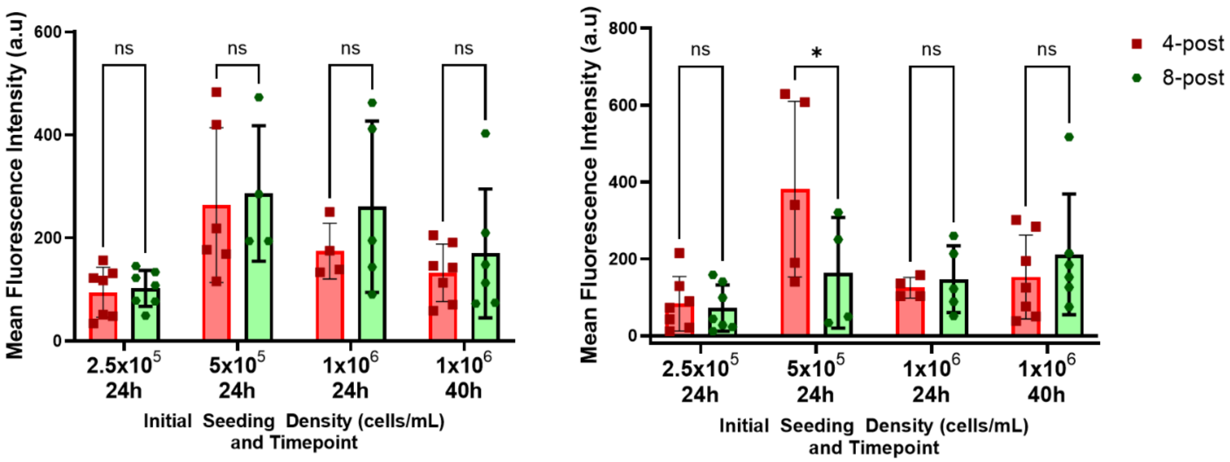


Figure S5. Analysis of the fluorescence intensity of α-SMA (left) and F-actin (right) staining within center region of 4-post and 8-post tissues at lowcell seeding densities (<2x10$^7$ cells/mL) at either 24 or 40 hours of culture. Each data-point represents one tissue. Error bars: *SD*. * $p < 0.05$, 2-way *t*-test. . Data from one batch representative of three biological replicates.

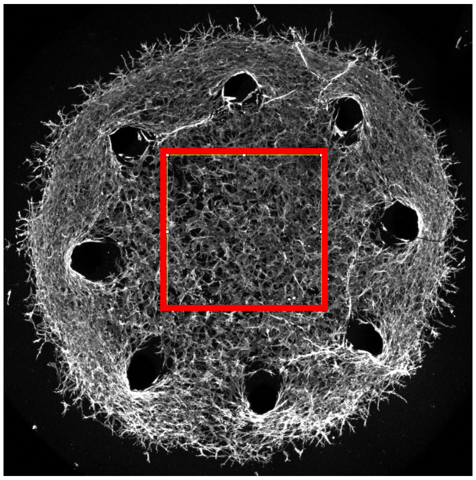

Figure S6. Example of 1 mm x 1 mm ROI selection at the center of the tissue specimen for cell and nuclei morphology analysis

**Supplementary Information on the Model**

The model predicts cell activation in response to the stress field in the tissue. Unlike thermoelastic-based classical cell models, our model accounts for how fibroblasts promote their actomyosin-based contractile forces (denoted by the average of $\rho_{kk}$) in response to both the magnitude of tension ($\sigma_{kk}$) and anisotropy of the tensile stress field ($\sigma_{\mathrm{a}}$) generated in the tissue (**Fig. 3a**). [1–3]

The model first predicts that for both 4-post and 8-post tissue designs, cell contractility and cell force generation increase with external resistance to contraction arising from increases in micro-post stiffness. These predictions are in agreement with experimental observations where the intensity of fibroblast activation biomarkers and tissue contractile force increased with micro-post stiffness.[1–4]

Second, the model predicts that cell contractility and cell force generation increase with anisotropy of the stress field and that the effect of tension anisotropy can overpower the effect of external resistance arising from increasing the number of micro-posts. Because the external resistance against tissue contraction in 8-post tissue is higher than in 4-post tissue, cells within the central region of the 8-post tissue are expected to experience higher tension and thus exhibit greater cell activation levels (**Fig. 3b)**. However, the model predicts that cells within the central region of the 4-post tissue, where the stress field is highly anisotropic, have higher actomyosin contractility and force generation levels compared with the cells within the central region of the 8-post tissue where the stress field is isotropic (**Fig. 3c,d).**[1–3] Similarly, the model predicts a similar level of cell activation around the periphery of 4-post and 8-post tissues as cells in those regions experience anisotropic stress fields in both tissues (**Fig. 3c,d**). Note that these predictions are opposite to the predictions from classical pre-stressed/pre-strained models, as these models do not account for the effect of tension anisotropy on cell activation and force generation levels (**Fig. 3b**).
